# Enhancing Percolation in Phosphatic Clay Using Diatoms under Laboratory Conditions

**DOI:** 10.1101/357889

**Authors:** Melba D. Horton, Dieff Vital, Paul Defino, Sarah Spaulding, Gary Albarelli

## Abstract

Clay settling areas are large impoundments brought about by phosphate mining with water-holding capacity that renders it very poor for agriculture and crop production. This study aims to enhance water percolation in phosphatic clay using porous diatom frustules under laboratory conditions. Phosphatic clay collected from settling areas was brought to the laboratory for the experiment. Diatom frustules were purchased commercially and dry samples of the diatom, *Didymosphenia* were provided by the University of Colorado-Boulder. Oven-dried clay was mixed with diatom frustules into 125 mL centrifuge tubes following a 1:1 volume ratio as experimental set-ups while pure phosphatic clay was used as control. Deionized water was poured into each set-up and the percentage of unpercolated water overlying the sediment, water retained in the sediment particles, water that percolated and passed through the hole of the centrifuge tube were monitored for 48 hours. Results showed that the addition of diatom frustules enhances the percolation of water in the sediment mixture especially those with *Didymosphenia* frustules. However, this mixture also showed higher percentage of water retained in the sediment particles which could be attributed to the high carbon and organic content brought about by the presence of stalks which is a major component of this species morphology. Considering how *Didymosphenia* disturbs freshwater habitats, proper management may render it useful for the mitigation of clay settling areas in the land environment. The implication of this on crop production remains to be explored and further *in situ* experimentations need to be conducted.

**IMPORTANCE:** Clay settling areas abound in places where phosphate mining is conducted. The very fine particles and the chemical property of this phosphatic clay allows it to hold water more than normal clay sediments making the area unstable and less suitable for agricultural use. Studies show that mitigative measures to enhance surface drainage is very costly leaving most areas barren and unused. Diatoms are unicellular algae in various size and shapes with silicified cell walls that are porous and are ubiquitous in aquatic environments. The significance of our research is being able to demonstrate the potential use of diatoms, most especially the genus *Didymosphenia* which is regarded as an environmental threat to some habitats, in mitigating the drainage problem in clay settling areas by mixing phosphatic clay with diatom frustules. This process is cost-effective and more importantly provides utilization of a resource that is regarded as nuisance in freshwater environments.

## INTRODUCTION

Percolation of water in the soil have different implications not only in terms of soil property (1), but also on the nutrients that could be possibly leached and become unavailable for plant use (2, 3). Macropores and matrix porosity add more complexity in trying to get a thorough understanding of how water flows in different soil types (4). Heavy clay soils reportedly exhibit more water logging due to low hydraulic conductivity (5, 6) brought about by particle-size distribution (7). Phosphatic clay is no exception and understanding its permeability is still a challenge (8).

The Institute of Food and Agricultural Science revealed that more than 100,000 acres of land in Florida have been dedicated to clay settling areas over the course of phosphate mining in Florida (9), with a large percentage of that area still remaining unreclaimed and not currently in use due to its poor quality making the settling areas unstable. A typical clay settling area covers up to 40% of the land for a given phosphate mine as suggested by the Florida Industrial and Phosphate Research Institute (http://www.fipr.state.fl.us/about-us/phosphate-primer/phosphogypsum-and-the-epa-ban/). Compared to other naturally-occurring soil types that are typically sandy and organic with less fertility and low water-holding capacity, phosphatic clay is man-made, very fertile and has a high water-holding capacity. Studies conducted on the potential of reclaimed phosphatic clay for crop production indicated low water infiltration rates that necessitates additional surface drainage which is very costly given the low marketability of the produce (10, 11).

Diatoms are photosynthetic single-celled algae with siliceous shells called frustules found in fresh and marine waters (12). Their ubiquitous distribution in the environment provides a significant impact on the carbon and silicon cycles. The unique properties of its porous frustules have been explored for a variety of applications to include ecological monitoring (13, 14), drug delivery, (15), heterogeneous catalysis (16), gene transfection carriage (17), and forensic research (12). Geological timeline indicates that diatoms have been depositing their siliceous skeletal remains in the sediment since the early Triassic period (18) as soft, granular, component called diatomaceous earth (19). During WWII, diatomaceous earth was reportedly used as a filtering agent to provide potable water, a technology that lasted until today (20). Diatomaceous earth is currently mined for commercial production and processed into two different grades; 1) the industrial grade is strictly used as a filtering agent in swimming pools with very high risk for human consumption, and 2) the food grade is reportedly safe for human consumption while serving as an excellent insecticide. This effect of the latter grade is also observed in pests and worms whereby the former can be lethally affected by the diatomaceous powder while the latter and other useful micro-organisms in the soil can stay unaffected (21, 22).

Considering the porosity of diatoms’ siliceous shells and its abundance in the environment, it is hypothesized that adding diatom frustules into the sediment mixture would be a cost-effective measure to lower the water-holding capacity of phosphatic clay to support crop production. This study is conducted to investigate the effect of adding diatom frustules into phosphatic clay in enhancing water percolation under laboratory conditions.

## MATERIALS AND METHODS

### Phosphatic Clay sample and analysis

Wet phosphatic clay were collected from settling areas by Mosaic’s South Fort Meade facility and transported by a FIPR representative to Florida Polytechnic University laboratory. Samples of the phosphatic clay were placed in petri dishes and oven-dried for 24 hours at 60°C. The dried samples were then powdered using mortar and pestle. One milligram of powdered sample was put on a stub for scanning electron microscopy (SEM) and energy-dispersive spectroscopic (EDS) analysis using an ultra-high-resolution Hitachi SU8200 Series.

### Diatom sample and analysis

Dried samples of the diatom, *Didymosphenia* were provided by Dr. Spaulding from the University of Colorado- Boulder. The bulk of stalks and frustules were powdered using mortar and pestle. Commercially available food grade diatomaceous earth was purchased. Both samples were examined and characterized using high resolution differential interference contrast (DIC) microscopy, SEM, and EDS.

### Phosphatic clay: Diatom Mixture preparation and data analysis

A 1:1 volume ratio of phosphatic clay: *Didymosphenia* and phosphatic clay: diatomaceous earth were weighed separately using a Mettler Toledo scale (SN B421637099) and each sediment mixture was vortex-mixed thoroughly for 10 minutes. Five milliliter of each mixture was poured into a 15 mL Falcon centrifuge tube with a hole (2.3 mm) drilled at the bottom tip to allow water flow. Each centrifuge tube was then inserted into a 25 mL graduated cylinder leaving a space between the bottom and the tip of the tube to collect any water outflow. Ten mL of deionized water was used to carefully pour into each centrifuge tube with the sediment mixture and the amount of water that 1) stayed overlying the sediment is considered unpercolated, 2) the amount that passes through the sediment through the hole of the centrifuge tube and is collected at the bottom of the cylinder is considered percolated, and 3) the difference between the percolated and unpercolated water volume is considered the amount retained within the sediment mixture.

Each sediment mixture; phosphatic clay:diatomaceous earth and phosphatic clay: *Didymosphenia* was prepared in 3 experimental replicates while 3 separate set-ups with only phosphatic clay served as control. Each set-up was monitored continuously for 48 hours and the amount of water that percolated, retained, and stayed unpercolated were recorded accordingly. Distribution of the data was tested for normality and the significance of the difference between samples was analyzed using analysis of variance (ANOVA) with the SPSS software package.

## RESULTS

### Characterization of phosphatic clay sample

EDS analysis shown in Fig. 1 revealed that phosphatic clay is by percent weight composed of oxygen (50.5), carbon (16.2), silicon (12.3), calcium (7.7), aluminum (5.4), magnesium (3.5), iron (2.3), phosphorus (1.5), and potassium (0.5).

**Fig. 1.**
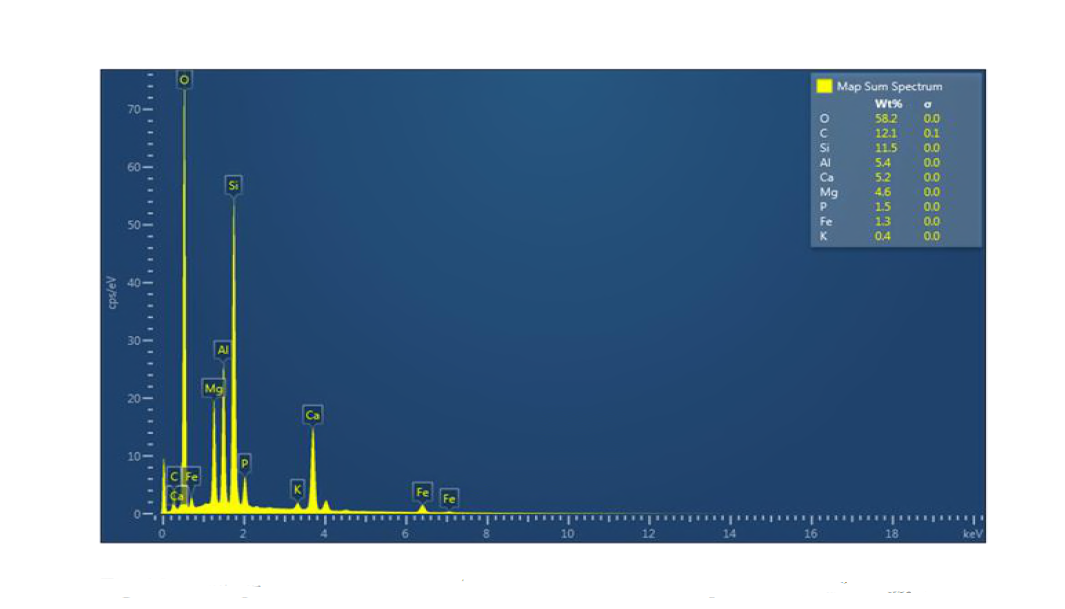
Percentage weight of the elemental composition of the phosphatic clay sample using energy dispersive spectroscopy (EDS).

### Characterization of diatom sample

Results of the DIC microscopy revealed that the diatomaceous earth sample is comprised of 100% diatom frustules in various shapes and sizes as shown in Fig. 2. EDS analysis showed that the diatomaceous earth is by percent weight composed of oxygen (57.1), carbon (2.7), silicon (32.5), calcium (0.7), aluminum (4.9), magnesium (0.8), iron (1.2), and titanium (0.2).

**Fig. 2.**
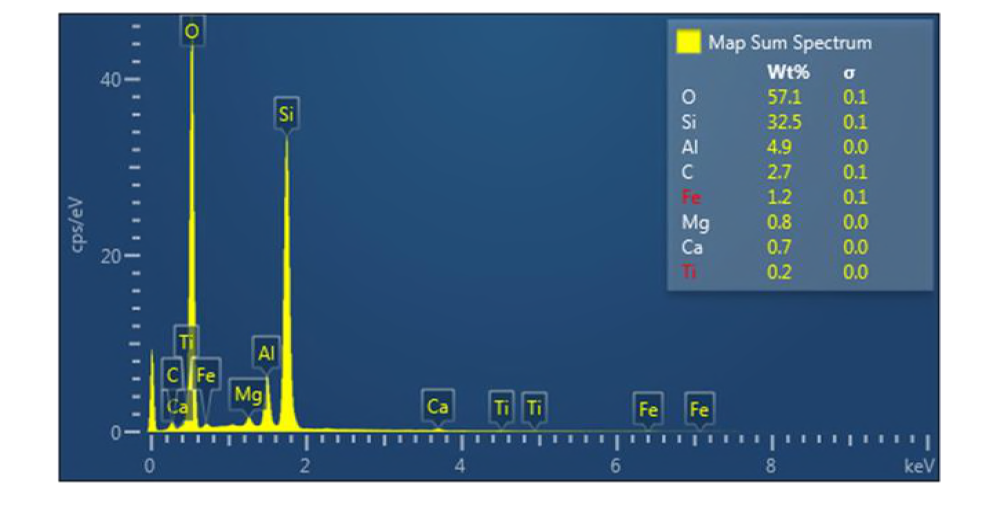
Percentage weight of the elemental composition of the diatomaceous earth sample using energy dispersive spectroscopy (EDS).

On the other hand, the *Didymosphenia* sample revealed the most number of elements identified in the EDS analysis as shown in Fig. 2c with the percentage by weight composition of oxygen (33.8), carbon (39.6), silicon (12.4), calcium (3.2), magnesium (1.3), iron (0.9), phosphorus (0.2), potassium (0.9), nitrogen, (5.7), sulfur (0.8), chlorine (0.7), and sodium (0.4) (Fig. 3).

**Fig. 3.**
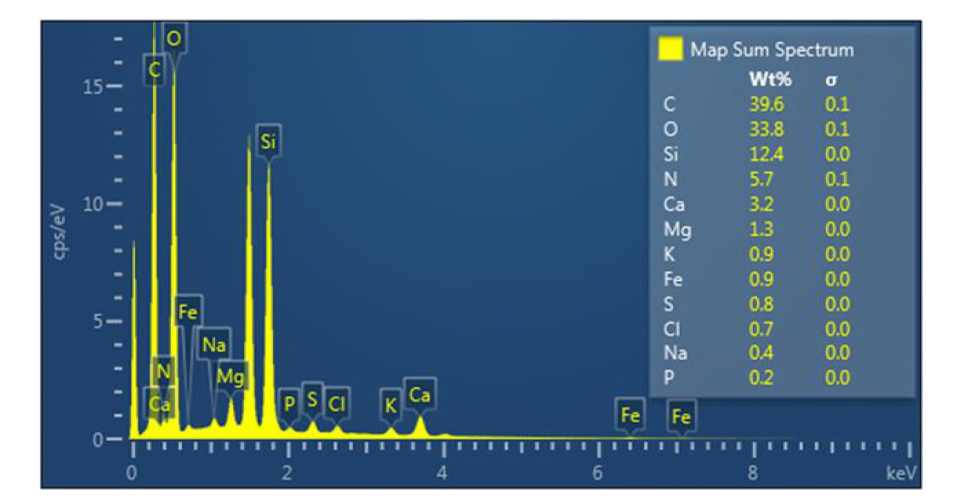
Percentage weight of the elemental composition of the *Didymosphenia* frustules and stalks sample using energy dispersive spectroscopy (EDS).

### Testing water percolation in sediment samples and statistical analysis

As shown in Fig. 4a, for the first 8 hours, about 70% of the water remained unpercolated in the phosphatic clay sediment sample, more than 20% was retained in the sediment particles and only less than 10% percolated which flowed through and out of the centrifuge tube. After 24 hours, the amount of unpercolated water showed a very minimal but steady decline to about 45% corresponded by an opposing trend of minimal increase in the small percentage that percolated to around 30% by the end of 48 hours. The amount of water retained in the phosphatic clay sediment became constant at about 25% after 32 hours. In the phosphatic clay:diatomaceous earth sediment mixture, the percentage of unpercolated water drastically dropped to less than 40% in the first 2 hours followed by a slow decline to less than 20% after 8 hours. More than 40% percolated through followed by a steady increase to about 55% and only about 25% was retained in the particles of the sediment mixture. After 28 hours, the percentage of water retained leveled off to less than 25% while the percentage that percolated continued to show minimal but steady increase to almost 70% with a corresponding decrease in the percentage of unpercolated water to about 5% at the end of 48 hours (Fig. 4b). In the phosphatic *clay:Didymosphenia* sediment mixture, the percentage of unpercolated water dropped logarithmically to about 10% after 8 hours. This logarithmic drop is coupled with the corresponding logarithmic increase in the percentage of percolated water to more than 60% while 30% was retained in the sediment mixture that remained constant from then on. After 24 hours, all the water (about 70%) that was not retained in the sediment mixture had already percolated (Fig. 4c).

**Fig. 4.**
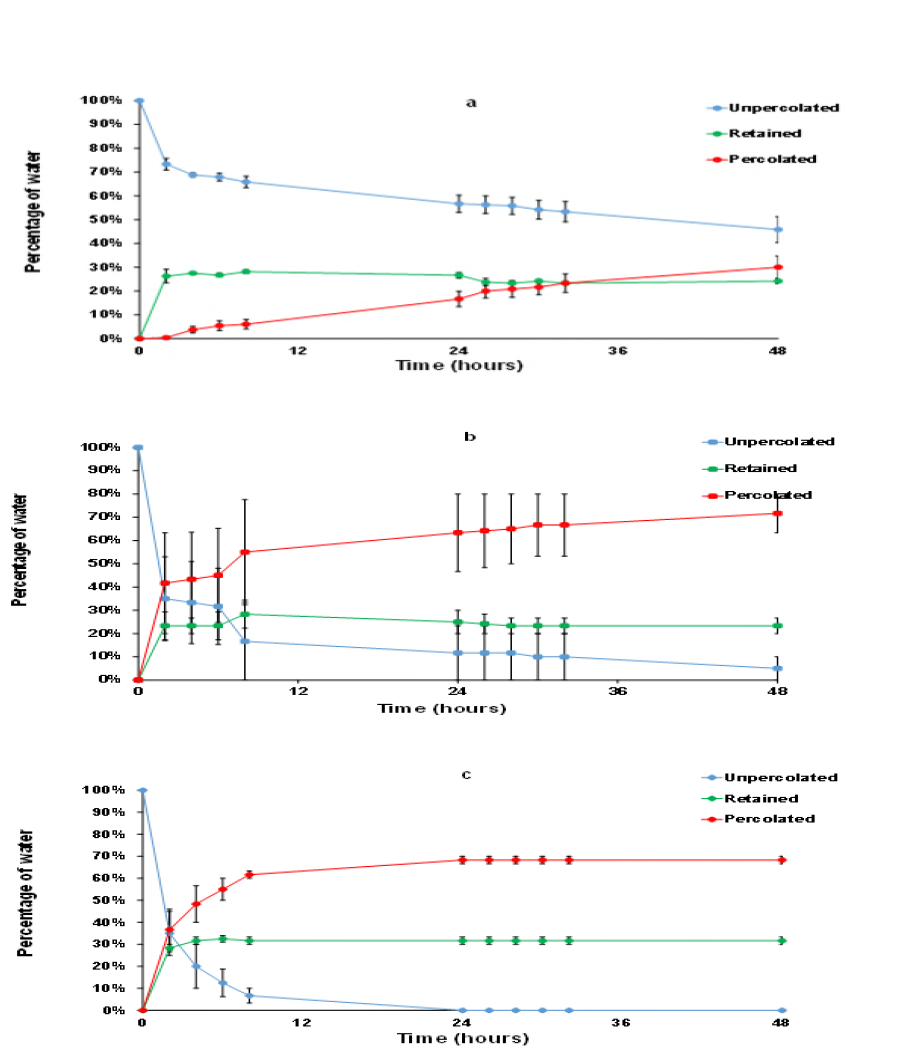
Average percentage of unpercolated water overlying the sediment, water retained in the sediment particles, and water that percolated through **a)** phosphatic clay (n=6; ±SE), **b)** phosphatic clay: diatomaceous earth (n=3; ±SE), and **c)** phosphatic clay: *Didymosphenia* (n=3; ±SE) frustules and stalks sediment mixtures in a 1:1 volume ratio under laboratory conditions.

Result of the analysis of variance (ANOVA) conducted on the difference in the amount of water that percolated, retained, and stayed unpercolated between and among the three sediment types tested revealed a significant difference at p≤ 0.05 level of significance as shown in Table 1.

**Table 1.** Table 1. Analysis of variance on the amount of water that percolated, retained, and stayed unpercolated among the the three types of sediments tested in a 1:1 ratio, namely: phosphatic clay:diatomaceous earth; phosphatic clay:*Didymosphenia*, and just phosphatic clay as control under laboratory conditions.

| Source of Variation | SS | df | MS | F | P-value |
| --- | --- | --- | --- | --- | --- |
| Between Groups | 594.45 | 3 | 198.15 | 83.35 | *8.34E-37 |
| Within Groups | 561.02 | 236 | 2.38 |  |  |
| Total | 1155.47 | 239 |  |  |  |

\* Significant at $p \leq 0.05$

## DISCUSSION

The development of terrestrial soils and the pore system that is formed in the process is the result of an interplay of the various subsystems of the planet’s spheres (23). Furthermore, the size of aggregate formation in clay dominated soil is influenced by the carbon content and other organic binding elements (24). As shown in the EDS results, phosphatic clay has low percentage of carbon as does the diatomaceous earth. This is probably why the amount of water retained in both soil samples is much lower than the one mixed with the diatom, *Didymosphenia.* As previously described, the sample comprised of stalks and frustules of this diatom. Gretz (25) reported that *Didymosphenia* stalks are composed of sulfated polysaccharides. This supports the high percentage of carbon as well as the presence of diverse elements shown in the EDS of this sample which possibly increased the binding of soil aggregates and in effect retained more water within the sediment mixture. Nonetheless, the phosphatic clay:*Didymosphenia* sediment mixture showed the fastest water percolation compared to the other sediment types. This could be attributed to the size of the highly porous diatom frustules of which *Didymosphenia* is reportedly larger than most species (26, 27). This might have influenced the particle size distribution in the clay particles, creating voids which may play an important role in water percolation (7). In contrast to the phosphatic clay: diatomaceous earth mixture, the impact is not readily observed which can be attributed to the smaller sizes of the frustules from a variety of diatom species that dominate the sample. This result provides an avenue to utilize *Didymosphenia*, which is regarded as a nuisance in freshwater environments (28, 29, 30) into a useful mitigative measure in clay setting areas with proper management plan.

The addition of diatom frustules in the phosphatic clay sediment in a 1:1 volume ratio clearly resulted in a significant increase in the percolation of water within a shorter period of time under laboratory condition. Utilizing the diatom, *Didymosphenia* in the mixture results in a faster percolation, however, the presence of stalks could also enhance the organic binding components which could result in a higher amount of water retained in the sediment particles.

The implication of this on plant growth and crop production remains to be explored, and how this results play *in situ*, which would require a tremendous amount of diatom sample, needs to be carried out for further investigation.

## ACKNOWLEDGEMENTS

This research was supported by the Florida Polytechnic University internal seed grant 85051603MH and the authors thank Florida Industrial of Phosphate Research Institute (FIPR) for generously providing the phosphatic clay for use in the experiments.

